# Diffusion Latent Representations for Neural Decoding

**DOI:** 10.64898/2026.07.08.737343

**Authors:** Brandon Wong, Brokoslaw Laschowski

## Abstract

Neural decoding can be viewed as a representation learning problem in which neural activity is mapped into an intermediate representation before downstream reconstruction. The choice of intermediate representation influences both performance and learning difficulty. Here we developed a novel framework for studying how intermediate representation choice influences downstream learning and reconstruction. As a proof-of-concept, we instantiated our framework using diffusion latent representations extracted from different diffusion timesteps for neural speech decoding. Component-wise evaluation showed that reconstruction performance differed substantially across diffusion timesteps, with teacher-forced Word Error Rates of 44.7%, 7.5%, and 3.5% for different latent models. These results demonstrate that diffusion latent representations can serve as effective intermediate representations for learning from neural activity, but that their effectiveness depends strongly on the selected diffusion timestep. More broadly, our framework provides a basis for systematically studying how intermediate representation choice influences downstream learning and reconstruction.

## I. Introduction

MACHINE learning systems rarely learn directly from raw inputs to downstream tasks. Instead, they transform data into intermediate representations that capture salient structure for subsequent prediction, generation, or decision-making. Representation learning has become a key paradigm in machine learning [1]–[3], underpinning advances in computer vision, natural language processing, speech recognition, reinforcement learning, and generative modeling. The choice of intermediate representation influences what information can be preserved, how efficiently it can be learned, and ultimately how effectively it supports downstream tasks. Understanding how representation choice influences learning remains an important problem in machine learning.

Neural decoding provides a natural framework for studying intermediate representations because its objective is to map neural activity onto representations that support downstream reconstruction. Neural activity is high-dimensional, noisy, and stochastic, making the choice of intermediate representation particularly important. Rather than decoding directly from neural activity to target outputs, many neural decoding algorithms first map neural activity into an intermediate representation before reconstruction by a downstream decoder. Previous research has explored a variety of intermediate representations for image decoding [4]–[6], speech decoding [7]–[10], and other neural decoding tasks [11]–[15]. Despite their differences, they all transform neural activity into intermediate representations that support downstream reconstruction. Collectively, these studies demonstrate that neural decoding can be viewed as a representation learning problem in which the choice of representation plays a central role in performance.

Intermediate representations possess distinct statistical and geometric properties that influence how effectively they can be learned from neural activity. Latent representations derived from generative models have emerged as a promising class of representations because they capture the underlying probability distribution of the target modality. For example, denoising diffusion probabilistic models [16] generate high-fidelity data by reversing a gradual noising process through a sequence of intermediate noisy variables. Diffusion-based representations have recently been applied to neural decoding, where latent diffusion models reconstruct visual stimuli from brain activity by mapping neural activity into the latent space of a pretrained diffusion model [4]. These advances motivate studying diffusion latent representations as a promising class of intermediate representations.

Despite growing interest in representations, relatively little attention has been paid to understanding how different intermediate representations influence learning from neural activity [17]. Previous studies have largely evaluated representations empirically through downstream performance, while the statistical and geometric properties that influence their learnability remain poorly understood. This limitation is particularly important in neural decoding, where high-dimensional and limited training data place strong demands on the choice of representation.

Motivated by this, here we developed a novel framework for studying how representation choice influences downstream learning and reconstruction. As a proof-of-concept, we instantiated our framework using diffusion latent representations extracted from different diffusion timesteps for neural speech decoding. Our framework predicts intermediate representations and conditioning factors directly from neural activity, enabling different representations to be systematically evaluated. More broadly, our framework provides a basis for studying how intermediate representation choice influences downstream learning and reconstruction.

## II. Methods

We developed this framework to study how representation choice influences downstream reconstruction. In this proof-of-concept implementation, our framework consisted of three stages: (1) generating diffusion latent representations and conditioning factors, (2) predicting them from neural activity using transformer encoder-decoders, and (3) reconstructing the target signal using a frozen diffusion model. Because our framework operates on intermediate representations, it is, in principle, adaptable to different conditional diffusion architectures.

### A. Encoder

The encoder autoregressively predicts the intermediate representations required by the diffusion decoder, which are tokenized diffusion latent representations and an associated conditioning factor. Our framework includes two transformer encoder-decoder models [18]: a latent model that predicts tokenized diffusion latents and a conditioning factor model that predicts conditioning factors from neural activity. Both models share the same transformer encoder backbone consisting of sinusoidal positional embeddings and multi-head self-attention. The latent model decoder jointly predicts all *J* Residual Vector Quantization (RVQ) codebook indices at each autoregressive step to account for the hierarchical structure of RVQ tokens. The conditioning factor model decoder uses a linear projection layer for continuous regression. Both models include a stop-probability head for autoregressive termination.

### B. Decoder

The decoder reconstructs the target stimulus from the tokenized diffusion latent representations and conditioning factor predicted by the encoder. Rather than learning to generate the target stimulus directly from neural activity, our framework uses a pretrained conditional diffusion model to exploit generative priors learned from large-scale datasets. To instantiate our framework, we used the conditional diffusion model DiffWave [19], although other conditional diffusion models could have been explored. In this study, the latent tokens were converted back to continuous latent variables using EnCodec, generating 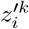 from 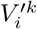. The decoder then performs reverse diffusion from the predicted latent variable according to

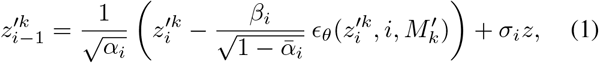

where *z* ∼ *N*(0, *I*) and *σ*_*i*_ is a DiffWave hyperparameter. The reverse diffusion process iteratively refines the latent variable from timestep *i* to the fully denoised state.

### C. Training

During training, the diffusion model remained frozen while only the encoder models were optimized. In this initial implementation, four transformer encoder-decoder models were trained: one conditioning factor model and three latent models corresponding to diffusion timesteps *i* ∈ {1, 2, 3}, denoted latent models 1–3, respectively. A separate latent model was trained for each timestep to avoid conditioning instability and to learn timestep-specific representations.

These timesteps were selected to balance signal detail and generative flexibility. We excluded *i* = 0 to avoid requiring the encoder to reconstruct fine-grained signal details directly from neural activity. Larger timesteps (*i* ≥ 4) were excluded because they contain less structural information, reducing the contribution of the latent representation while placing greater emphasis on the conditioning factor.

All models were trained using teacher forcing with a hidden size of 128, 4 transformer layers, and 2 attention heads. The optimization used Adam with a learning rate of 5 × 10^−4^, batch size 128, and 100 training epochs. No learning-rate scheduler or dropout was used. We optimized the latent models using the summed cross-entropy loss across all *J* Residual Vector Quantization codebooks, while the conditioning factor model used Mean Squared Error loss. The stop-probability heads of both models were optimized using binary cross-entropy. All models were implemented in PyTorch and trained using DistributedDataParallel on four NVIDIA H100 GPUs.

### D. Dataset

As a representative application, we evaluated our framework on a speech decoding dataset [20] consisting of intracortical recordings from a participant implanted with a 256-electrode microelectrode array. Neural activity was binned into non-overlapping 20 ms windows containing threshold crossings and spike band-power features, yielding 512-dimensional feature vectors at each timestep. We used the predefined training and validation splits consisting of 7,879 and 1,409 neural activity– sentence pairs, respectively.

### E. Training Target Generation

To instantiate our framework, we first generated the intermediate representations used as training targets for the encoder models. These targets consisted of diffusion latent representations and conditioning factors required by a pretrained conditional diffusion model. The selected conditioning factor depends on the underlying diffusion model. For example, conditional diffusion models may use action labels for motion generation [21] or text prompts for image generation [22]. In this initial implementation, we used DiffWave [19], which conditions on Mel spectrograms. Ground-truth audio waveforms were synthesized from the target sentences using PiperTTS with the `US-amy-medium` voice to generate the conditioning factors.

We computed Mel spectrograms from the synthesized audio and performed conditional reverse diffusion using DiffWave to obtain diffusion latent representations. Latent representations were extracted at every diffusion timestep (*T* = 6 for DiffWave). Because the resulting latent trajectories are prohibitively long for autoregressive prediction, we tokenized each latent representation using EnCodec [23], which applies Residual Vector Quantization across *J* codebooks. Here, *J* = 4 with 1024 entries per codebook. We first generated the Mel spectrograms, denoted 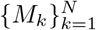, where *N* is the total number of target sentences. Each Mel spectrogram can be represented as the sequence

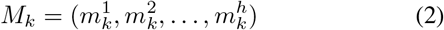

where *h* is the sequence length and each vector has dimension 80. We then generated the corresponding diffusion latent representations by performing conditional reverse diffusion. The resulting latent representations are denoted 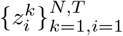, where *k* indexes the sample and *i* denotes the diffusion timestep, *i* ∈ {1, 2, …, *T*}. After tokenization, each latent representation 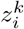 is represented as

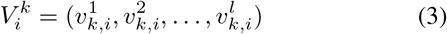

where each token belongs to the product space of the RVQ codebooks,

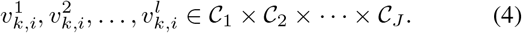

### F. Inference and Evaluation

During inference, both encoder models autoregressively generate the intermediate representations required by the diffusion decoder from a single beginning-of-sequence (BOS) token. The conditioning factor model predicts the conditioning factor together with a stop probability used to determine the output sequence length. In this initial demonstration, the conditioning factor is a Mel spectrogram. The optimal stop threshold was selected from the range [0.1, 1.0] using the validation set. Given the predicted Mel spectrogram length *L*_*mel*_, the latent sequence length *L*_*lat*_ was computed by aligning the temporal resolutions of the Mel spectrogram and EnCodec latent space,

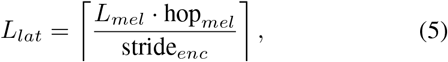

where hop*mel* is the Mel spectrogram hop length and stride_*enc*_ is the EnCodec encoder stride. The latent model then autoregressively predicts a latent sequence of length *L*_*lat*_. The predicted latent tokens are decoded using EnCodec and combined with the predicted conditioning factor before being passed to the frozen diffusion model for reconstruction. We evaluated our framework using two complementary analyses: (i) end-to-end evaluation of the complete framework and (ii) component-wise evaluation of the individual encoder models, such that for the conditioning factor model, the predicted conditioning factor was provided to the diffusion model.

For the latent models, the predicted latent representations were injected at their corresponding diffusion timesteps while conditioning on the ground-truth conditioning factor. This evaluation isolated each representation from errors introduced by the other encoder model, allowing their contributions to downstream reconstruction to be assessed independently. The end-to-end evaluation was performed under both teacher-forced and fully autoregressive generation, whereas component-wise evaluation isolated each encoder model under teacher forcing. Synthesized audio was resampled to 16 kHz, transcribed using Whisper [24], and evaluated using Word Error Rate (WER),

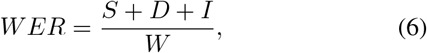

where *W* denotes the number of ground-truth words, and *S, D*, and *I* are substitutions, deletions, and insertions, respectively. Our source code is available here.

## III. Results

Using our framework, we first evaluated how representation choice influenced downstream reconstruction. Table I shows the component-wise Word Error Rate (WER) when each encoder model was evaluated under teacher-forced generation. This analysis isolates the contribution of each intermediate representation. We found that performance differed substantially across diffusion timesteps. The conditioning factor model achieved a WER of 3.5%, while the latent models achieved 44.7%, 7.5%, and 3.5% for latent models 1–3, respectively. Latent model 3 matched the performance of the conditioning factor model, whereas earlier diffusion latent representations produced much higher reconstruction errors. These results demonstrate that representation choice strongly influences downstream learning and reconstruction.

**TABLE I.**
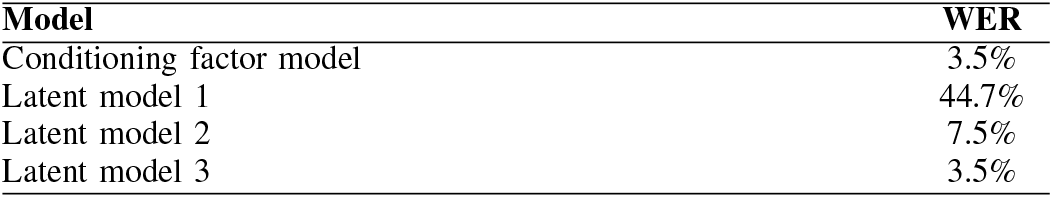
Teacher-forced Word Error Rate (WER) for independently evaluated models at the final training epoch.

| Model | WER |
| --- | --- |
| Conditioning factor model | 3.5% |
| Latent model 1 | 44.7% |
| Latent model 2 | 7.5% |
| Latent model 3 | 3.5% |

Figure 2 summarizes the optimization of the encoder models throughout training. The top row shows the training and validation losses and the bottom row shows the component-wise evaluation of each encoder model throughout training. All models showed stable optimization with converging loss curves. However, downstream reconstruction performance differed substantially across the diffusion timesteps. For example, latent model 3 consistently achieved the lowest WER throughout training, whereas latent models 1 and 2 converged to higher reconstruction errors despite similarly stable optimization. In contrast, the conditioning factor model demonstrated low WER throughout training. These results indicate that low training loss alone does not guarantee an effective intermediate representation for downstream reconstruction.

**Fig. 1.**
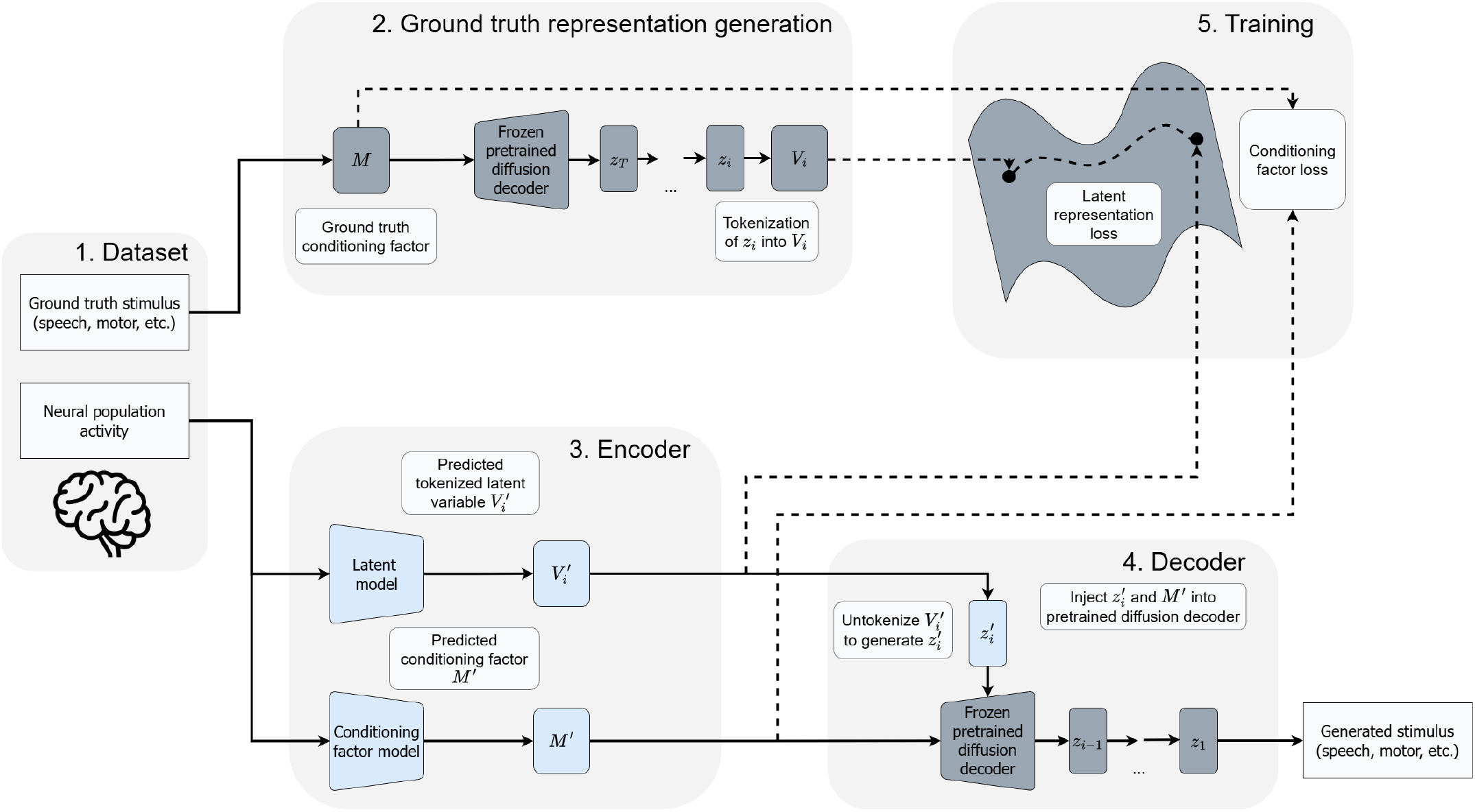
Overview of our framework for studying how representation choice influences downstream learning and reconstruction. A pretrained diffusion model performs reverse diffusion using the ground-truth conditioning factor *M* to generate target latent variables *z*_*i*_, which are tokenized into *V*_*i*_. The latent model maps neural activity onto tokenized latent representations, which are then untokenized into 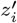. Simultaneously, the conditioning factor model predicts *M*′ using the ground-truth conditioning factor *M* as the target. During inference, 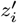 and *M*′ are injected into the pretrained diffusion model to reconstruct the target stimulus.

**Fig. 2.**
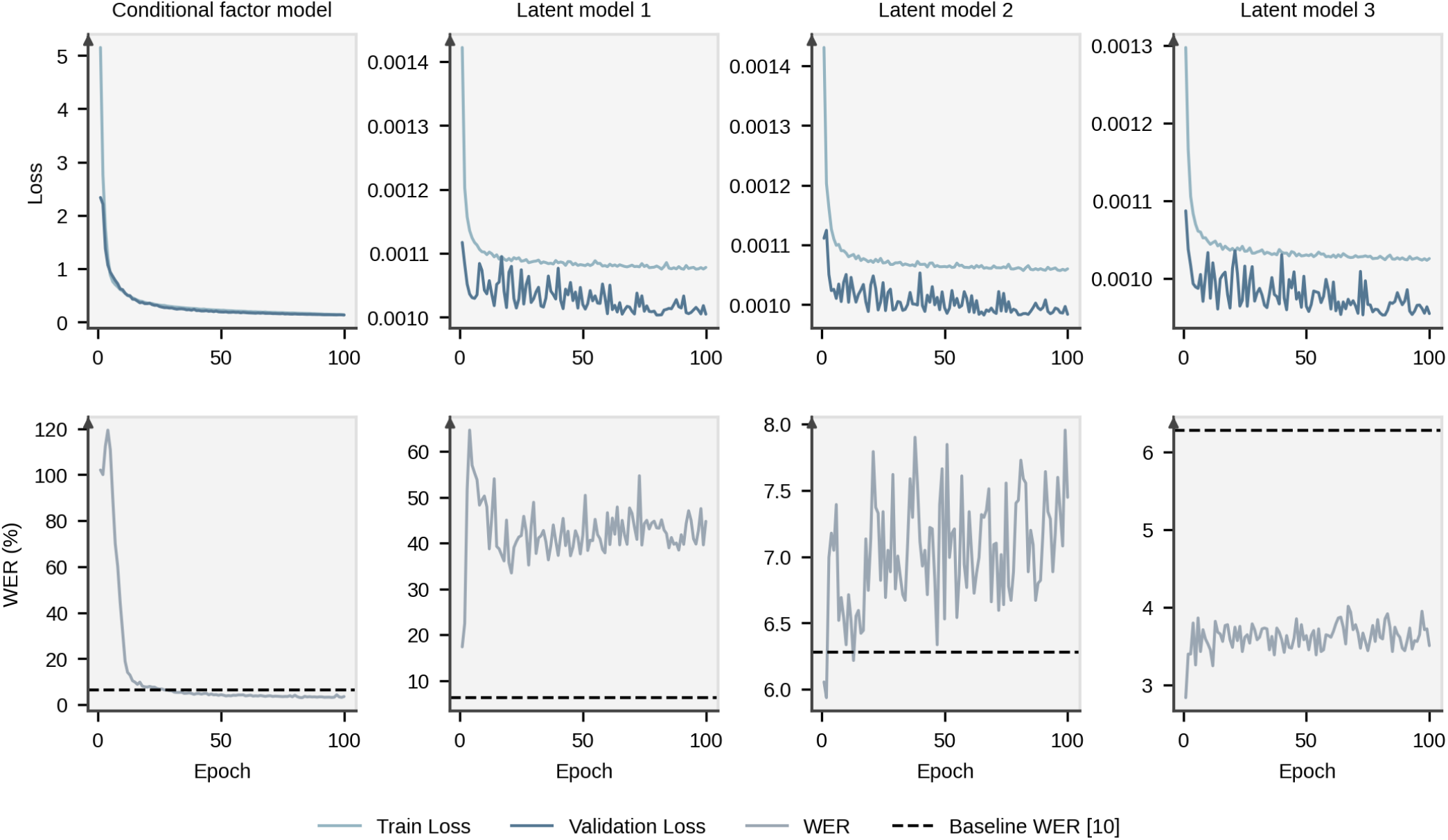
Training dynamics for the conditioning factor and latent models. The top row shows training and validation losses and the bottom row shows validation Word Error Rate (WER) under teacher-forced generation. Although all models showed stable optimization, downstream reconstruction performance depended on the intermediate representation.

We also evaluated the complete framework by combining the conditioning factor model with the highest-performing latent model, latent model 3. Table II compares our framework with previous research [10] on the same validation dataset. Under teacher-forced generation, our complete framework achieved a WER of 4.6%, demonstrating that the selected diffusion latent representation can support accurate downstream reconstruction when combined with a pretrained conditional diffusion model. In contrast, fully autoregressive generation yielded a WER of 125.3%, revealing that autoregressive error accumulation remains a major limitation of our current implementation. This discrepancy suggests that performance degradation mainly arises during autoregressive sequence generation rather than reconstruction from the predicted intermediate representations.

**TABLE II.**
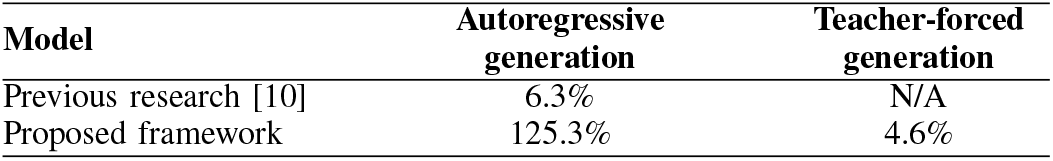
Comparison with previous research using autoregressive and teacher-forced generation.

| Model | Autoregressive generation | Teacher-forced generation |
| --- | --- | --- |
| Previous research [10] | 6.3% | N/A |
| Proposed framework | 125.3% | 4.6% |

## IV. Discussion

In this study, we developed a novel framework for studying how representation choice influences downstream learning and reconstruction. As a proof-of-concept, we instantiated our framework using diffusion latent representations extracted from different diffusion timesteps for neural speech decoding. Using this framework, we found that reconstruction performance differed substantially across diffusion timesteps, demonstrating that representation choice can strongly influence downstream reconstruction performance. Although this initial implementation focused on diffusion latent representations, the proposed framework is designed to be applicable to a broad range of intermediate representations and neural decoding tasks, providing a basis for systematically studying how representation choice influences downstream learning and reconstruction.

Our main finding is that representation choice strongly influenced downstream reconstruction performance. Previous neural decoding studies have shown that intermediate representations can support accurate reconstruction across a variety of tasks [4], [7], [9]–[11]. However, these studies mainly focused on a single intermediate representation selected for a specific application. In contrast, our method isolates the effect of intermediate representation choice by systematically comparing different representations within a controlled downstream reconstruction framework. For example, latent model 3 achieved a teacher-forced WER of 3.5%, whereas latent models 2 and 1 achieved 7.5% and 44.7%, respectively. One possible explanation is that different diffusion timesteps correspond to regions of the latent space with distinct statistical and geometric properties. Later diffusion timesteps may preserve greater stochasticity and allow the diffusion model to iteratively correct reconstruction errors during subsequent denoising steps, whereas earlier diffusion timesteps may require increasingly precise latent predictions directly from neural activity. Although not directly evaluated, this interpretation is consistent with the training dynamics shown in Figure 2, where earlier diffusion timesteps exhibited less stable WER convergence despite relatively stable optimization loss.

Another important finding of our research is that the complete framework performed well under teacher-forced evaluation (4.6% WER) but poorly under autoregressive generation (125.3% WER). Similar to previous neural decoding research [9], [10], our proof-of-concept implementation relied on autoregressive sequence generation. Our results suggest that errors introduced during sequential prediction accumulated over time, substantially degrading end-to-end performance despite accurate intermediate representations.

Looking ahead, future research should extend this framework to compare representations derived from different generative and representation-learning models to determine which statistical and geometric properties best facilitate learning from neural activity. Our findings build on the growing literature on representation learning [1]–[3] by suggesting that representation choice should be evaluated not only by downstream reconstruction performance, but also by how effectively representations can be learned from neural activity. Beyond neural decoding, this research contributes toward the broader aim of studying representations learned by biological and artificial neural systems [25]–[27], bridging the gap between computational neuroscience and machine learning.

## V. Conclusion

Here we developed a novel framework for studying how intermediate representation choice influences downstream learning and reconstruction. As a proof-of-concept, we instantiated our framework using diffusion latent representations extracted from different diffusion timesteps for neural speech decoding. Component-wise evaluation demonstrated that reconstruction performance differed substantially across diffusion timesteps, highlighting that representation choice can strongly influence downstream reconstruction performance. More broadly, the proposed framework provides a foundation for comparing intermediate representations across neural decoding tasks and contributes toward a deeper understanding of representation learning in biological and artificial neural systems.

## Acknowledgments

This research is dedicated to the students and researchers in Ukraine. Their resilience and unwavering commitment to education and learning continue to serve as a beacon of hope and inspiration to the global academic community.

## Notes

This research was partially supported by the Schroeder Institute for Brain Innovation and Recovery.

### Competing Interest Statement

The authors have declared no competing interest.

## eferences

[1] Y. Bengio, A. Courville, and P. Vincent, “Representation learning: A review and new perspectives,” IEEE Transactions on Pattern Analysis and Machine Intelligence, vol. 35, no. 8, pp. 1798–1828, 2013.

[2] I. Goodfellow, Y. Bengio, and A. Courville, Deep Learning. MIT Press, 2016.

[3] T. Chen, S. Kornblith, M. Norouzi, and G. Hinton, “A simple framework for contrastive learning of visual representations,” in International Conference on Machine Learning (ICML), 2020, pp. 1597–1607.

[4] Y. Takagi and S. Nishimoto, “High-resolution image reconstruction with latent diffusion models from human brain activity,” in IEEE/CVF Conference on Computer Vision and Pattern Recognition (CVPR), 2023, pp. 14 453–14 463.

[5] Z. Guo, J. Wu, Y. Song, J. Bu, W. Mai, Q. Zheng, W. Ouyang, and C. Song, “Neuro-3d: Towards 3d visual decoding from eeg signals,” in IEEE/CVF Conference on Computer Vision and Pattern Recognition (CVPR), 2025, pp. 23 870–23 880.

[6] K. N. Kay, T. Naselaris, R. J. Prenger, and J. L. Gallant, “Identifying natural images from human brain activity,” Nature, vol. 452, no. 7185, pp. 352–355, 2008.

[7] S. d’Ascoli, C. Bel, J. Rapin, H. Banville, Y. Benchetrit, C. Pallier, and J.-R. King, “Towards decoding individual words from non-invasive brain recordings,” Nature Communications, vol. 16, no. 1, p. 10521, 2025.

[8] G. K. Anumanchipalli, J. Chartier, and E. F. Chang, “Speech synthesis from neural decoding of spoken sentences,” Nature, vol. 568, no. 7753, pp. 493–498, 2019.

[9] A. Défossez, C. Caucheteux, J. Rapin, O. Kabeli, and J.-R. King, “Decoding speech perception from non-invasive brain recordings,” Nature Machine Intelligence, vol. 5, no. 10, pp. 1097–1107, 2023.

[10] N. S. Card, M. Wairagkar, C. Iacobacci, X. Hou et al., “An accurate and rapidly calibrating speech neuroprosthesis,” New England Journal of Medicine, vol. 391, no. 7, pp. 609–618, 2024.

[11] A. Dempster and B. Laschowski, “Mixture models for domain-adaptive brain decoding,” bioRxiv, 2025.

[12] A. Motiwala, J. Soldado-Magraner, A. P. Batista, M. A. Smith, and B. M. Yu, “Brain–computer interfaces as a causal probe for scientific inquiry,” Trends in Cognitive Sciences, vol. 30, no. 1, pp. 40–53, 2026.

[13] A. G. Kurbis, A. Mihailidis, and B. Laschowski, “An emg foundation model for neural decoding,” bioRxiv, 2025.

[14] O. Shevchenko, S. Yeremeieva, and B. Laschowski, “Comparative analysis of neural decoding algorithms for brain-machine interfaces,” in IEEE International Conference on Rehabilitation Robotics (ICORR), 2025, pp. 222–227.

[15] J. P. Cunningham and B. M. Yu, “Dimensionality reduction for large-scale neural recordings,” Nature neuroscience, vol. 17, no. 11, pp. 1500–1509, 2014.

[16] J. Ho, A. Jain, and P. Abbeel, “Denoising diffusion probabilistic models,” in Advances in Neural Information Processing Systems (NeurIPS), vol. 33, 2020, pp. 6840–6851.

[17] M. Fuest, P. Ma, M. Gui, J. Schusterbauer, V. T. Hu, and B. Ommer, “Diffusion models and representation learning: A survey,” IEEE Transactions on Pattern Analysis and Machine Intelligence, 2026.

[18] A. Vaswani, N. Shazeer, N. Parmar, J. Uszkoreit, L. Jones, A. N. Gomez, Ł. Kaiser, and I. Polosukhin, “Attention is all you need,” Advances in Neural Information Processing Systems (NeurIPS), vol. 30, 2017.

[19] Z. Kong, W. Ping, J. Huang, K. Zhao, and B. Catanzaro, “Diffwave: A versatile diffusion model for audio synthesis,” arXiv preprint arXiv:2009.09761, 2020.

[20] N. S. Card, M. Wairagkar, C. Iacobacci, X. Hou et al., “Brain-to-text ‘25,” https://kaggle.com/competitions/brain-to-text-25, 2025, kaggle.

[21] G. Tevet, S. Raab, B. Gordon, Y. Shafir, D. Cohen-Or, and A. H. Bermano, “Human motion diffusion model,” arXiv preprint arXiv:2209.14916, 2022.

[22] A. Ramesh, P. Dhariwal, A. Nichol, C. Chu, and M. Chen, “Hierarchical text-conditional image generation with clip latents,” arXiv preprint arXiv:2204.06125, 2022.

[23] A. Défossez, J. Copet, G. Synnaeve, and Y. Adi, “High fidelity neural audio compression,” arXiv preprint arXiv:2210.13438, 2022.

[24] A. Radford, J. W. Kim, T. Xu, G. Brockman, C. McLeavey, and Sutskever, “Robust speech recognition via large-scale weak supervision,” in International Conference on Machine Learning (ICML). PMLR, 2023, pp. 28 492–28 518.

[25] S. Kostousov, A. Kaushik, and B. Laschowski, “Bidirectional representational alignment between biological and artificial neural networks,” bioRxiv, 2026.

[26] S. Leno, A. Petrovych, and B. Laschowski, “Topological regularization for neural representational alignment,” bioRxiv, 2026.

[27] B. A. Richards, T. P. Lillicrap, P. Beaudoin, Y. Bengio et al., “A deep learning framework for neuroscience,” Nature Neuroscience, vol. 22, no. 11, pp. 1761–1770, 2019.

